# Enhancing Opioid Bioactivity Predictions through Integration of Ligand-Based and Structure-Based Drug Discovery Strategies with Transfer and Deep Learning Techniques

**DOI:** 10.1101/2023.08.04.552065

**Authors:** Davide Provasi, Marta Filizola

## Abstract

The opioid epidemic has cast a shadow over public health, necessitating immediate action to address its devastating consequences. To effectively combat this crisis, it is crucial to discover better opioid drugs with reduced addiction potential. Artificial intelligence-based and other machine learning tools, particularly deep learning models, have garnered significant attention in recent years for their potential to advance drug discovery. However, utilizing these tools poses challenges, especially when training samples are insufficient to achieve adequate prediction performance. In this study, we investigate the effectiveness of transfer learning using combined ligand-based and structure-based molecular descriptors from the entire opioid receptor (OR) subfamily in building robust deep learning models for enhanced bioactivity prediction of opioid ligands at each individual OR subtype. Our studies hold the potential to greatly advance opioid research by enabling the rapid identification of novel chemical probes with specific bioactivities, which can aid in the study of receptor function and contribute to the future development of improved opioid therapeutics.

## 1. INTRODUCTION

The opioid epidemic has led to an urgent need for the development of safer and more effective opioid drugs. While various strategies, such as the development of opioid receptor (OR) allosteric modulators, biased agonists, and compounds with low efficacy are being pursued in both academic and industry settings, the development of opioids with reduced side effects and the ability to effectively treat opioid use disorders remain a significant challenge.^1, 2^

To expedite and improve the drug discovery process, computer-aided drug discovery techniques can be invaluable.^3^ There are two main approaches in computer-aided drug discovery: ligand-based (LB) and structure-based (SB) methods. LB methods can help in the design of novel opioid ligands by predicting their biological activity based on shared molecular descriptors with known opioids. On the other hand, SB methods involve utilizing three-dimensional protein structures, specifically those of δ-, μ-, and κ-opioid receptors (DOR, MOR, and KOR) in this case, to design and optimize opioid molecules. By examining the atomic-level interactions between opioids and receptors, computational models can be employed to identify potential chemical modifications of known opioids or new compounds that could enhance the desired therapeutic effects while minimizing side effects.

Although both LB and SB methods have made significant contributions to drug discovery^4^ and machine learning (ML) has been widely used in LB virtual screening, the integration of these approaches with artificial intelligence (AI)-based techniques is still in its infancy when it comes to applications to members of the G Protein-Coupled Receptor (GPCR) family, such as ORs. However, this integration holds the promise of taking the field to a new level. AI-based tools, particularly deep learning models, can analyze vast amounts of data, identify patterns, and make accurate predictions.^5^ By training AI models on large databases of chemical and biological information, one can expedite the discovery of compounds with desired pharmacological profiles, potentially reducing the current lengthy drug discovery timeline and associated exorbitant costs. Some of these methods have proven successful in applications aimed at accelerating GPCR drug discovery, including the prediction of GPCR structures, ligand-GPCR interactions, clinical responses, and novel GPCR ligands, whether targeting orthosteric or allosteric sites.^6–8^ In these applications, chemical or structural descriptors have typically served as inputs for classification models that employ support vector machines, random forest, naïve Bayes classifiers, and neural networks.^9–11^

Despite the recent successful applications, it is important to note that not all GPCRs are well-suited for AI-based tools, particularly for deep learning applications, due to the limited availability of structural and functional data for some of them. To overcome this limitation, transfer learning, a powerful technique in ML, can also be used to pre-train models on large-scale datasets and subsequently fine-tune these models on specific datasets.^12^ This fine-tuning allows for a quick training and good performance of the model even if training data from the specific dataset is limited. In recent applications to GPCRs, transfer learning allowed a more accurate modeling of ligand-GPCR associations when combined with molecular graphs convolutional networks.^13^

Here, we investigate an integrated approach that harnesses the synergistic information provided by LB and SB techniques along with transfer learning to improve the accuracy of opioid bioactivity prediction across the three OR subtypes, using a dense neural network (DNN)^14^ or graph convolutional neural network (GCN)^15^ architecture. While the use of hybrid LB+SB computational schemes has shown improvements in virtual screening results and the overall success of drug discovery endeavors,^4^ to the best of our knowledge, this is the first time that information pertaining to both ligands and GPCR structures has been integrated within a transfer learning framework for bioactivity predictions.

## 2. COMPUTATIONAL METHODS

### 2.1 Biological Activity Information for Training and Testing Datasets

Training datasets of biologically active (agonists or antagonists) ligands at each of the three main subtypes of the OR subfamily (DOR, MOR, or KOR dataset) or the combined dataset (OR subfamily dataset, excluding the nociceptin/orphanin FQ opioid peptide receptor) were retrieved from the IUPHAR/BPS Guide to Pharmacology (Release 2023.1)^16^, whereas corresponding datasets of inactive ligands were obtained from the ChEMBL database (Release 32, 2023-01-26)^17, 18^. Large linear peptides, identified as molecules with a molecular weight greater than 510 Da, with at least one match for substructure queries for amino acids or a dipeptide group, having more than 10 rotatable bonds, and with the largest ring containing fewer than 11 atoms, were removed from these datasets. The remaining ligands annotated in the Guide to Pharmacology database as agonists or antagonists at DOR, MOR, and KOR (codes 317, 319, and 318, respectively) or reported in the ChEMBL database with-log_10_ potency values (-log_10_Ki, –log_10_IC50, or –log_10_EC50) lower than 5 (or Ki, IC50, and EC50 > 10 μM) at either human DOR, MOR, or KOR (codes 236, 233, and 237) were considered active or inactive ligands, respectively, for the purpose of training DNN and GCN models. Compounds with opposing bioactivity data reported in ChEMBL were excluded from the datasets. Table S1 provides the total number of ligands in the respective datasets, as well as their mean potency values. An additional ChEMBL dataset of “potent active” opioid ligands (-log_10_Ki, –log_10_IC50, or –log_10_EC50 > 7 or Ki, IC50, or EC50 < 10 nM), excluding compounds with reported opposing bioactivities, was prepared and used as a test set. Table S2 lists the total number of these “potent active” opioid ligands from ChEMBL, as well as their mean potency values.

### 2.2 Experimental Structural Information for Training Datasets

The Protein Data Bank (PDB) identification codes for all available ligand-OR complexes were obtained by querying the RCSB PDB Search API.^19^ The search was performed using the opioid receptor InterPro identifier (IPR001418) in the RCSB polymer entity annotation and requiring a non-polymer entity count greater than 0 in the RCSB entry info. Entries corresponding to the nociceptin/orphanin FQ opioid peptide receptor were manually excluded. At the time of this work, the search returned experimental structures for 25 complexes (3 DOR, 14 MOR, and 8 KOR). The corresponding PDB identification codes are listed in Table S3.

### 2.3 LB and SB Molecular Descriptors

The SMILES strings of all ligands were converted to canonical form using the CanonSmiles function from rdKit.^20^ A total of 43 LB molecular descriptors encompassing basic physical and chemical properties of the molecule, atom counts, and topological indices (see details in Table S4) were calculated using rdKit’s ComputeProperties function. SB molecular descriptors were calculated using iChem’s cavity-based pharmacophores.^21^ The strategy for calculating these descriptors consisted of (i) Applying the Schrödinger 2020-4 protein preparation workflow^22^ to the coordinates of each ligand-OR complex (see Table S3), (ii) Processing the resulting structures in mol2 format with the iChem Volsite tool^21^ in ligand-restricted mode to derive pharmacophoric features from the cavity around the ligand, (iii) Sampling probable tautomeric forms and protonation microspecies at pH 7.4, generating up to 200 three-dimensional (3D) conformers for each molecule, after enumerating stereoisomers for up to three stereocenters using the Quacpac 2.1.3.0^23^ and Omega 4.1.2.0^24^ toolkits from OpenEye 2021.2.0^25^, and (iv) Calculating the similarity of each of the conformers to each cavity-derived pharmacophore with the Shaper2 tool^21^ using default parameters. The SB molecular descriptors for each ligand were derived from the maximum, average, and 75% quantile of the distribution of the similarity values. By including various statistics related to the distribution of scores for an ensemble of ligand conformations, instead of just focusing on the maximum score, we could capture basic measures of the ligand’s flexibility in each binding pocket. Specifically, for each ligand *i* and each of the 25 structures μ, we considered three features corresponding to the maximum score, the average score, and the 75th percentile of the scores over the ensemble *C*(*i*) of microspecies and conformations for the ligand:

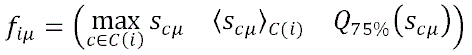

As a result, the dimensions of the SB feature vector for each ligand were 25×3=75.

### 2.4 Neural Network Architectures for Classification

Two distinct neural network architectures, specifically a DNN and a GCN, were used for classification. These followed concepts described in the literature^14, 15^ and are illustrated in Figures 1 and 2, respectively. The sizes of the relevant layers were chosen after hyperparameter optimization (see section 2.5) and are indicated in red in Figures 1 and 2. For the DNN model, the 43 LB and/or 75 SB molecular descriptors described in the previous section were used as input to two parallel functions: (i) A stack of two dense layers with Rectified Linear Unit (ReLU) activation, each followed by a dropout layer; and (ii) A single dense layer with ReLU activation, followed by a dropout layer. The output vectors of these two branches were concatenated and supplied to a final dense layer, which produced class probabilities via a soft-max activation function.

**Figure 1.**
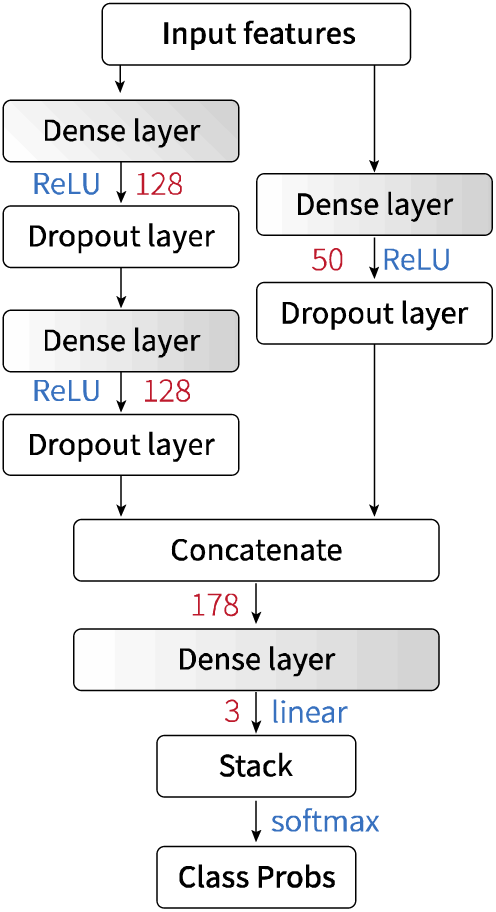
Architecture of the DNN classifier. Input and output dimensions of the layers are indicated in red when not obvious, while layers with trainable parameters are shaded in gray. Rectified linear unit (ReLU), linear, and soft maximum (softmax) activation functions are denoted in blue.

**Figure 2.**
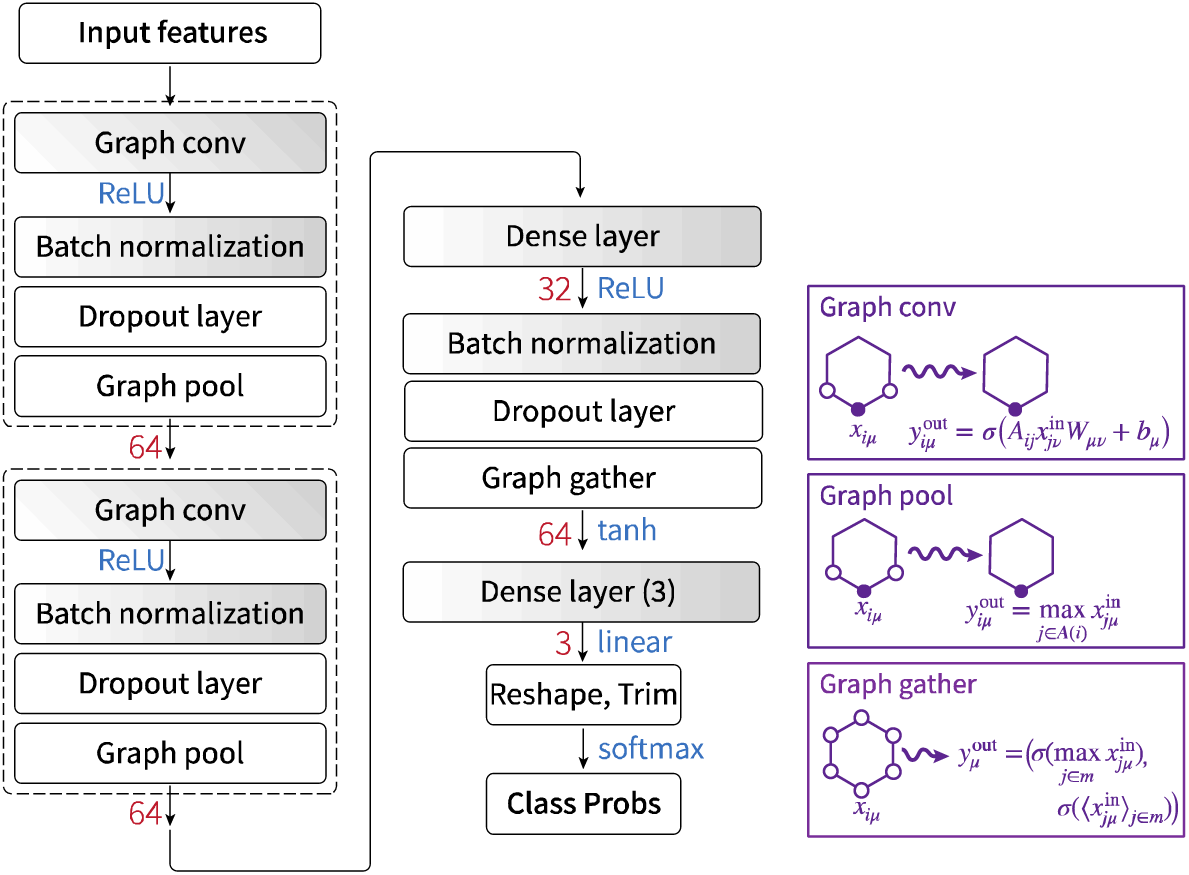
Architecture of the GCN classifier. Input and output dimensions of the layers are indicated in red, while layers with trainable parameters are shaded in gray. Rectified linear unit (ReLU), hyperbolic tangent (tanh), linear, and soft maximum (soft-max) activation functions are denoted in blue. The effect of the graph layers is described in the purple boxes, where Latin indices enumerate atoms, Greek indices enumerate atomic features, and summation over repeated indices is intended.

The GCN model was implemented using the DeepChem open source package.^26^ Initially, as per the default representation used in DeepChem, each atom was characterized by calculating a 75-long vector composed of: (i) A one-hot encoding of the atom’s name from a list of 44 possible values; (ii) A one-hot encoding of the number of atoms directly bonded to it (from 0 to 10); (iii) A one-hot encoding of the atom’s implicit valence (as calculated by rdKit’s GetImplicitValence) ranging from 0 to 7; (iv) The atom’s formal charge and the number of radical electrons (from rdKit’s GetFormalCharge and GetNumRadicalElectrons); (v) A one-hot encoding of the atom’s hybridization state (from the list sp, sp2, sp5, sp3d, sp3d2); (vi) A flag indicating if the atom is aromatic; and (vii) A one-hot encoding of the total number of implicit hydrogens on the atom (as calculated by GetTotalNumHs) ranging from 0 to 4. These atomic-level features were analyzed by two convolutional blocks and by a dense layer that produced atom-based embeddings. Subsequently, a gathering layer generated a molecule embedding, which was transformed by a hyperbolic tangent activation function and further analyzed by a final dense layer to yield class probabilities. The convolutional blocks were comprised of a GraphConv layer, followed by a batch normalization and a GraphPool layer. The GraphConv layer combined node (i.e., atom) feature vectors with the feature vectors of neighboring nodes. Specifically, using Latin indices to indicate atoms and Greek indices to indicate features, the output feature μ for atom *i* can be defined as:

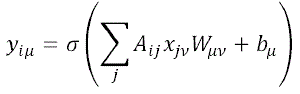

where x represents the input features, A is the normalized adjacency matrix, W is a learned kernel, and *b*, is a bias. This layer was followed by a batch normalization and a drop-out layer, and then by a Graph max-pool layer.^15^ Together with the convolution, this allowed the information in the local neighborhood of each atom to be combined as:

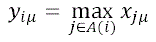

where *A*(*i*) indicates the atoms connected to *i*. This entire convolutional block was then followed by a second block and a dense layer, which generated atom-based embeddings as:

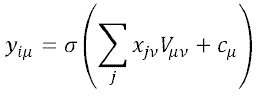

where *V* is a fully connected learned kernel and *c* is a bias. Finally, after another normalization and drop-out layer, the node-specific embeddings were transformed into a molecule-specific embedding by a graph gather layer:

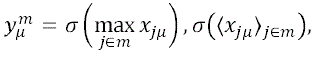

This layer calculated the maximum and average of each atomic features across the whole molecule and then applied the hyperbolic tangent activation function σ to transform it.

### 2.5 Training Protocols

An Adam optimizer (with β1=0.9, β2=0.999, ε=10^-8^, and a learning rate of 0.001) was used to minimize the SoftmaxCrossEntropy loss function. The minibatch size was set to 100. In the GCN models, the momentum for the batch normalization layers was set to 0.99. All training datasets were balanced with sample weights, ensuring equal total weights for each class in each of the datasets. Both SB and LB molecular descriptors were pre-processed before training by applying a centering transformation. The training datasets were divided into 10-fold cross-validation splits, with 90% allocated for training and 10% for testing. For transfer learning, after optimization on the pre-training training set, the parameters of all layers except the last dense layer of each model were fixed for an initial 50-epoch optimization on the fine-tuning training set. After this step, these parameters were released and further optimized for 50 epochs. Hyperparameters such as the number of nodes in the hidden layers, the number of nodes in the by-pass layer, and the drop-out probability, as well as the number of training epochs were optimized by training the models on the whole opioid dataset.

Statistics for different values of the hyperparameters are reported in Figures S1, S2, and S3 for the DNN classifiers and in Figure S4 for the GCN models. After inspecting the performance values, a common architecture was chosen for all the DNN models. This architecture had hidden layers of 128 nodes, a bypass layer with 50 nodes, a dropout rate of 0.2, and 100 training epochs. The GCN models used layers of 64 nodes in the convolutional blocks, dense layers of 32 nodes, and drop-out rates of 0.2 for all drop-out layers.

### 2.6. Saliency Estimation

Salient features in the dense architectures were identified by calculating the gradient of the final probability corresponding to a predicted class. If *m*(*f*)ɛ Δ^2^ represent the output probabilities in the standard 3-dimensional simplex Δ^2^ assigned by the trained model *m* to the three classes (agonist, antagonist, and inactive compounds) for a given input feature *f* and if *k* = argmax*_i_ m_i_*(*f*) represents the predicted class, the gradient can be calculated as follows:

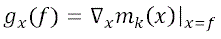

and the saliency can be measured by normalizing the absolute values component-wise:

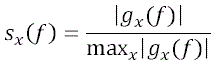

Values of *s_x_* close to 1 indicate that the feature *x* is maximally responsible for the prediction for the ligands with features *f* while values close to 0 indicate that the model does not use the corresponding features for the classification. We calculated the saliency for each feature in all correctly predicted ligands.

### 2.7 Performance statistics

Precision and recall, along with other metrics used to evaluate classifier performance, rely on the definition of a relevant class. Recall, for example, measures the fraction of correctly classified cases out of the total relevant cases (e.g., active compounds). In the studied setup, two out of the three classes can be naturally identified as “relevant” classes, namely the agonists and the antagonists. Therefore, we calculate the statistics for each class separately using a one-vs-rest approach and report the average of the two values for the active classes in Figures 3 and 4. The mean, 5^th^-percentile, 95^th^-percentile, and their averages over 10-fold samples of the class-specific values are also reported in Tables S5-S8.

**Figure 3.**
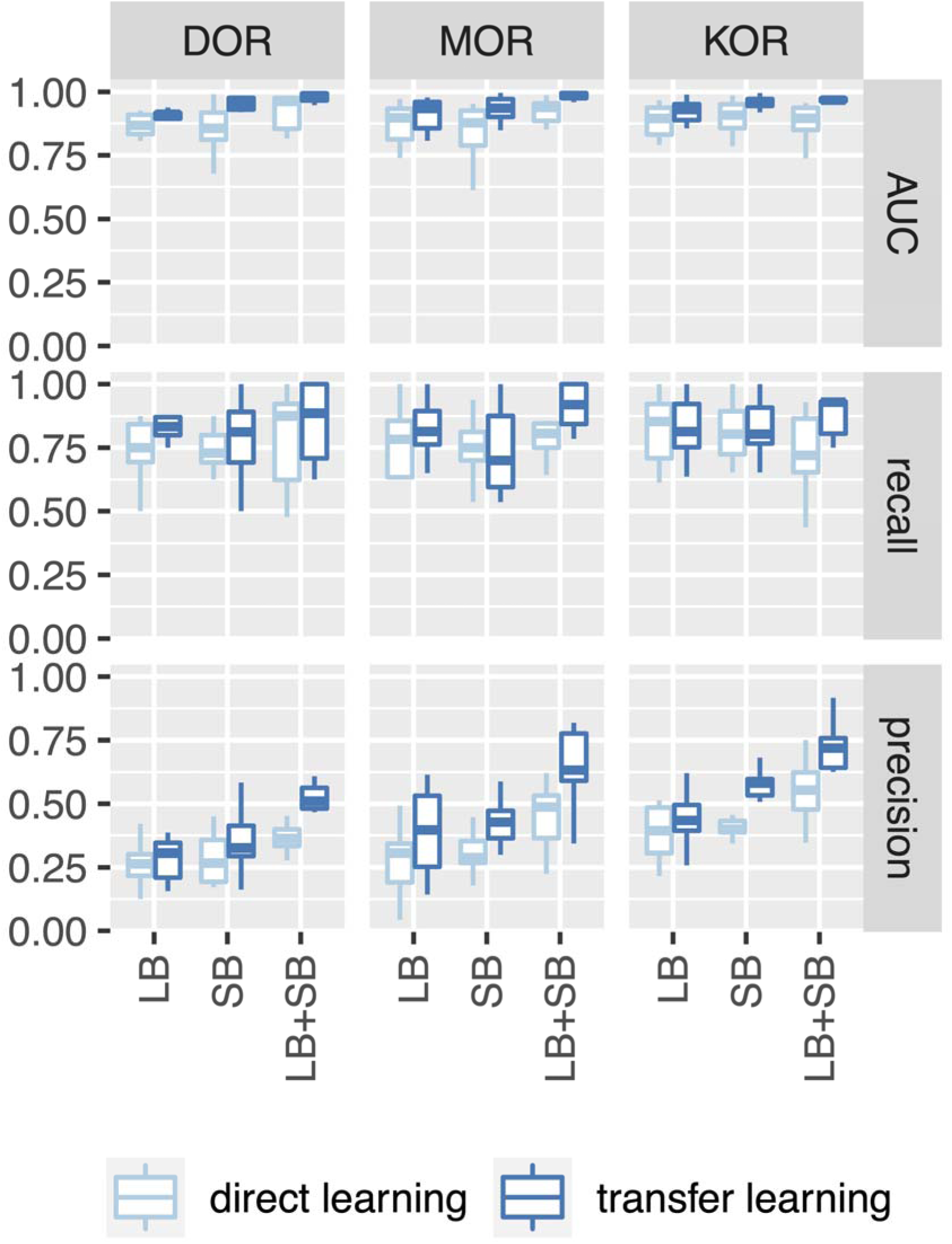
Performance metrics for the DNN classifiers. The boxplots represent the distribution of performance values on the test sets obtained from a 10-fold split of the training data. The metrics are calculated separately for each class, and the averages of the agonist and antagonist classes are plotted. AUC refers to the area under the Receiver Operating Characteristic (ROC) curve. LB, SB, and LB+SB refer to the models trained on ligand-based, structure-based, and combined features, respectively. Light blue and dark blue boxes represent direct and transfer learning values, respectively.

**Figure 4.**
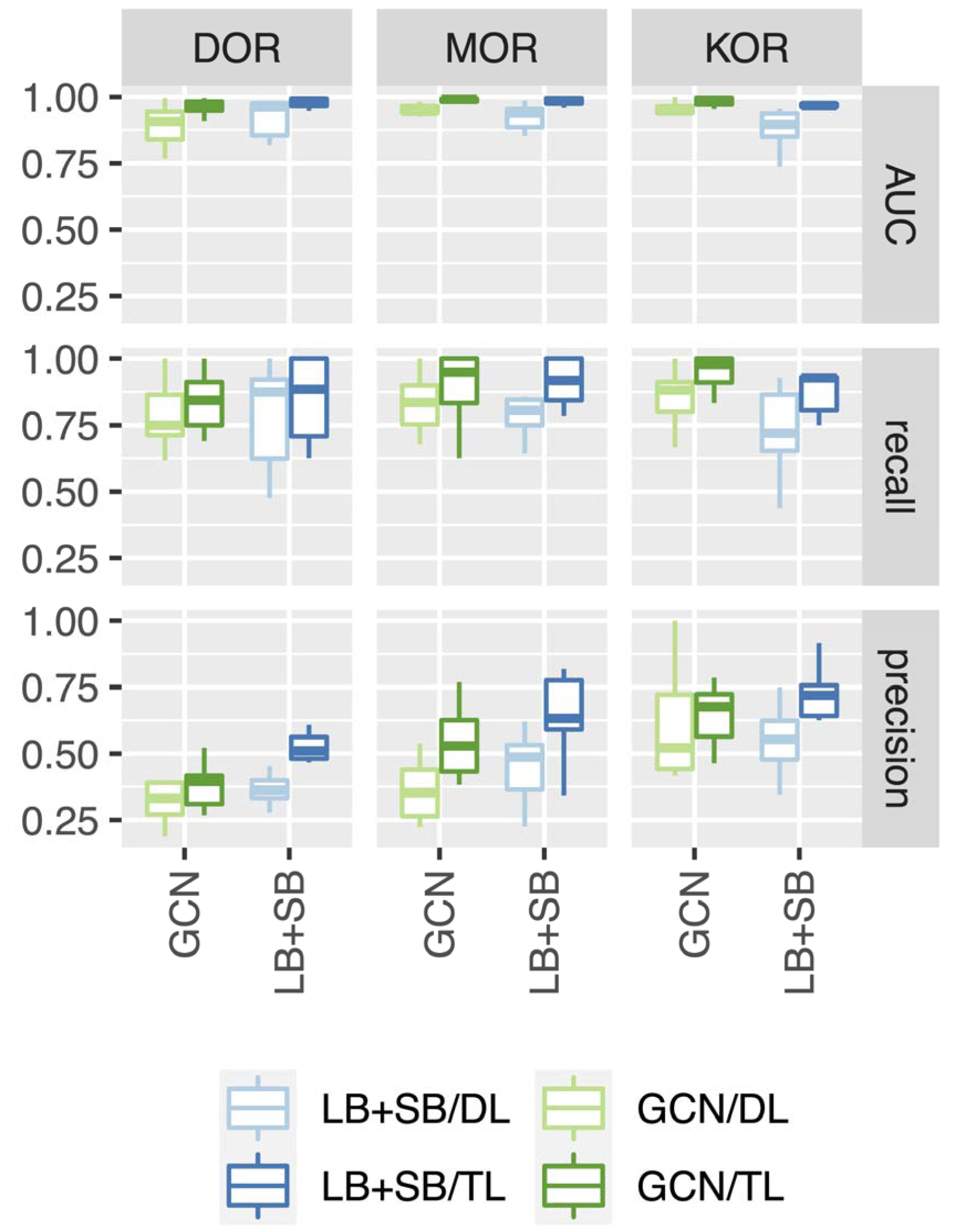
Comparison of the performance metrics between the DNN classifier trained on the combined LB+SB features (blue) and the LB-only GCN classifier (green). The boxplots represent the distribution of performance values on the test sets derived from a 10-fold split of the training data. The metrics are calculated separately for each class, and the averages of the agonist and antagonist classes are plotted. Light and dark blue colors represent results from training using direct learning (DL) and transfer learning (TL) protocols, respectively.

## 3. RESULTS AND DISCUSSION

With the overarching objective of enhancing the prediction of opioid bioactivity at individual ORs, and notwithstanding the limited ligand and structural information available, we evaluated the effectiveness of integrating LB and SB drug discovery strategies with transfer learning and two different deep learning models. To begin, we assessed the performance of DNN classifiers that employ a series of densely connected layers to process LB and SB molecular descriptors and estimate the functional properties of a molecule. For LB molecular descriptors, we selected a set of 43 physical and chemical properties, atom counts, and topological indices (see details in Table S4). Given the significant role of molecular recognition in ligand bioactivity, we considered SB molecular descriptors that quantified a molecule’s ability to fit the binding pocket of existing experimental OR structures. Specifically, these SB molecular descriptors comprised 75 measurements of the structural alignment between a ligand and cavity-derived pharmacophores of the binding pocket within the available 25 ligand-bound OR structures at the time of the study (see details in the Methods section and Table S3). The performance of DNN classifiers in distinguishing between agonist, antagonist, and inactive ligands of DOR, MOR, and KOR (see Methods section and Table S1) was first assessed by learning directly from either the 75 SB molecular descriptors or the 43 LB molecular descriptors independently, or from a combination of the two sets.

Results of the statistical analysis across 10-fold test splits are reported in Figure 3, and in Tables S5, S6, and S7 for DOR, MOR, and KOR, respectively. Figure S5 provides the corresponding performance metrics for the training sets. Although simple DNN models trained using either LB or SB molecular descriptors of individual ORs exhibit good area under the ROC Curve (AUC) scores (average values of 0.87, 0.87, and 0.88 for DNN models using LB features, and 0.86, 0.85, and 0.90 for DNN models using SB features for DOR, MOR, and KOR, respectively; see Tables S5, S6, and S7) and good recall (average values of 0.75, 0.69, and 0.82 for DNN models using LB features, and 0.76, 0.75, and 0.82 for DNN models using SB features for DOR, MOR, and KOR, respectively), their precision (average values of 0.27, 0.27, and 0.39 for DNN models using LB features and 0.28, 0.30, and 0.40 for DNN models using SB features for DOR, MOR, and KOR, respectively) is insufficient for practical application. By employing combined LB+SB descriptors in a DNN model trained on individual ORs, there is a marginal improvement in precision, reaching approximately 40% (average values of 0.37, 0.45, and 0.55, for DOR, MOR, and KOR respectively). However, this improvement remains relatively low, indicating a higher occurrence of false positives in these models. Upon examining the precision of the DNN models for each class in Tables S5, S6 and S7, it becomes evident that the agonist class exhibits lower precision compared to the antagonist class. This suggests a higher proportion of inactive or antagonist ligands being mistakenly predicted as agonists. On the other hand, the high precision achieved in the prediction of the inactive compounds may be a direct consequence of the corresponding dataset being much larger compared to the agonist and antagonist datasets.

Notably, adopting a transfer learning training protocol improves the precision of the DNN models up to 60% on average (specifically, average values of 0.51, 0.64, and 0.73, for the models using LB+SB features for DOR, MOR, and KOR, respectively). This improvement indicates that transfer learning effectively reduces the false discovery rate. Moreover, this reduction is accompanied by an overall increase in recall, reaching an average of 88% (average values of 0.85, 0.91, and 0.88 for DOR, MOR, and KOR, respectively) for the DNN models trained with transfer learning using combined LB+SB features. Therefore, the LB+SB combination not only reduces the likelihood of misclassifying an inactive ligand as active (precision) but also enhances the true positive rate (recall), which represents the fraction of accurately classified active ligands among all real active ligands.

To elucidate the important elements used by the classifiers to achieve their performance, we computed the saliency of each input feature in the trained models for correctly predicting ligands in those models. Saliency values enable the identification of the most influential descriptors contributing to the final classification of a given input molecule. The salient features for each of the fitted models in the three subfamilies, namely DOR, MOR, and KOR, are presented in Figures S6, S7, and S8, respectively. In general, the salient features identified in the DNN models trained with direct or transfer learning using LB or SB features are also salient in the DNN models trained with direct or transfer learning using combined LB+SB features, indicating that the combined models learn from the same features as the individual models. Several salient LB properties are common among all ORs. Specifically, properties such as Lipinski’s hydrogen-bond donor character,^27^ namely the sum of NH and OH bonds (lipinskiHBD), the number of hydrogen-bond donors (NumHBD), the number of saturated rings (NumSaturatedRings), and the number of atoms shared between rings that share exactly one atom (NumSpiroAtoms), all have average saliency values larger than 0.5 for all ORs, suggesting that these molecular features are important for a ligand’s bioactivity at opioid receptors.

Several SB properties of the ligand (defined by their complementarity to receptor’s binding pockets) with saliency values larger than 0.5 are also common among all OR models. These properties correspond to best, average, or 75^th^-percentile of similarity scores to cavity-derived pharmacophores in antagonist-bound DOR (PDB code 4N6H), agonist-bound MOR (PDB codes 8EF6 and 7U2L), and agonist– and antagonist-bound KOR structures (PDB codes 8DZR and 6VI4, respectively). Notably, the salient SB properties for the classification of ligands at an individual OR are not limited to those learned from structures of that OR but also rely on the structures of the other receptor subtypes. For instance, the salient SB properties used to classify DOR bioactive ligands are defined not only by DOR structures (PDB codes: 4EJ4 and 4N6H) but also by MOR (PDB codes: 7UL4, 7U2L, 8EF6, and 8EFL) and KOR structures (PDB codes: 6VI4 and 8DZR) (Figure S6). Similarly, MOR bioactive ligands are defined not only by MOR structures (PDB codes: 4DKL, 7SCG, 7T2H, 7U2L, and 8EF6) but also DOR (PDB code: 4N6H) and KOR structures (PDB codes: 6VI4 and 8DZR) (Figure S7). Finally, KOR bioactive ligands are defined by DOR (PDB code: 4N6H), MOR (PDB codes: 7UL4, 5C1M, 7U2K, 7U2L, 8EF6, and 8EFL), as well as by KOR (PDB codes: 6VI4, 6B73, 8DZQ, and 8DZR) structures (Figure S8). These observations suggest that DNN models do not simply use the best structural similarity between a ligand and known bioactive ligands for a specific OR as a basis for classification but rather rely on structural features defined also by other OR subtypes.

To illustrate the practical impact of the improvement, we identified active OR ligands that were consistently misclassified by directly trained DNN models using LB features but were correctly classified by DNN models using LB+SB features trained using transfer learning. Representative examples are shown in Figure S9. Interestingly, some morphinan ligands were predicted in the wrong active class by the simpler DNN models. For instance, nalbuphine, a non-selective agonist (partial agonist at DOR), was misclassified as an antagonist by DNN models trained through direct learning of LB features in 75% and 100% of the cases for DOR and KOR, respectively. However, it was always correctly classified by DNN models trained through transfer learning of combined LB+SB features (Figure S9, panels A and C). Similarly, nalfurafine, a selective full agonist at KOR, was misclassified as antagonist by DNN models trained through direct learning of LB features but was always correctly classified by DNN models trained through transfer learning of combined LB+SB features (Figure S9, panel C). BU72, a high-affinity MOR full agonist, was misclassified as an antagonist in 55% of the cases by DNN models trained through direct learning of LB features but was always correctly predicted by DNN models trained through transfer learning of combined LB+SB features (Figure S9, panel B). Transfer learning models that combine structural and chemical information can also correctly predict some active ligands with atypical chemical scaffolds that are consistently predicted to be inactive by the simpler DNN models trained by direct learning of LB features. Among these, BW373U86, a DOR agonist that also activates KOR with lower potency, was misclassified as inactive at KOR in 60% of the cases by DNN models trained through direct learning of LB features (Figure S9, panel C); etonitazene, a non-selective benzimidazole opioid (partial agonist at KOR), was always classified as inactive at MOR and KOR by DNN models trained through direct learning of LB features (Figure S9, panels B and C). Both ligands were correctly classified as inactive by DNN models trained through transfer learning of combined LB+SB features (Figure S9, panels B and C).

Next, we aimed to compare the performance of DNN classifiers to that of GCN classifiers, with both being trained using both the direct learning and transfer learning protocols. The results of the statistical analysis on 10-fold test splits, averaged across the two active classes, are reported in Figure 4 and are compared to the corresponding DNN models using combined LB+SB features. Averages and class-specific values are reported in Table S8 for each OR subtype. We note that transfer learning is advantageous for the GCN architecture as well. Across all metrics, and for each OR subtype, the models trained with transfer learning consistently outperformed those trained with direct learning.

The performance of the LB-only GCN architecture exceeds that of the DNN models using LB features. Specifically, when focusing on models trained with transfer learning, the use of a GCN model improves the AUC score (from average values of 0.91 to 0.96 for DOR, 0.91 to 0.98 for MOR, and 0.92 to 0.98 for KOR), recall (from average values of 0.82 to 0.84 DOR, 0.79 to 0.9 for MOR, and 0.82 to 0.95 for KOR), and precision (from average values of 0.28 to 0.38 for DOR, 0.39 to 0.55 for MOR, and 0.44 to 0.64 for KOR), as detailed in Table S8 and compared to Tables S5-S7.

In terms of AUC scores and recall, the LB-only GCN model performs comparably to the DNN model using LB+SB features. The AUC scores of the LB+SB model align with those of the LB-only GCN model (0.98 vs 0.96 for DOR, 0.98 vs. 0.98 for MOR, and 0.97 vs. 0.98 for KOR), as do most of the recall values (0.85 vs. 0.84 for DOR, 0.91 vs. 0.90 for MOR, and 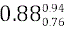 vs. 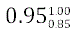 for KOR), as shown in Figure 4 and Tables S5-S8. However, it is worth noting that the precision achieved with a DNN classifier trained on the combined LB+SB descriptors still surpasses that of a LB-only GCN architecture. The precision of the LB-only GCN models decreases from the LB+SB value of 0.51 to 0.38 for DOR, from 0.64 to 0.55 for MOR, and from 0.73 to 0.64 for KOR (Figure 4 and Tables S5-S8). Therefore, we can conclude that while a GCN architecture can extract more relevant information from the 2D chemical structure of a ligand compared to a simpler DNN model using LB features, the integration of structural data remains beneficial for enhancing the precision of the classification.

To gain further insight into model’s performance on a more diverse dataset, we explored whether the models trained to separate agonists, antagonists, and inactive ligands could also be used to classify active vs. inactive ligands without distinguishing their specific efficacy. Specifically, we applied the trained classifiers to potent active and inactive opioid ligands extracted from ChEMBL, categorizing each ligand as active if the sum of the predicted probabilities for belonging to the active classes for each OR exceeded the predicted probability for being classified as an inactive compound at that receptor. The precision and recall values for this classification are reported in Table S9 and shown in Figure 5. First, we observe that, even for this simpler classification task on an extended dataset, transfer learning improved the performance (both precision and recall) of the trained DNN and GCN models. Secondly, for all subfamilies, the DNN models trained with either direct or transfer learning using LB features show very high recall values (e.g., 0.90, 0.91, and 0.81 for models trained with transfer learning for DOR, MOR, and KOR, respectively) and moderate precision (e.g., 0.58, 0.73, and 0.76 for models trained with transfer learning for DOR, MOR, and KOR, respectively), indicating that these models tend to classify some inactive molecules as active. Combining LB+SB features yields DNN models with better precision using either direct learning (0.72, 0.75, and 0.86 for DOR, MOR, and KOR, respectively) or transfer learning (0.81, 0.87, and 0.91 for DOR, MOR, and KOR, respectively) for training, but at the cost of reduced recall for models using both direct learning (0.74, 0.56, and 0.71 for DOR, MOR, and KOR, respectively) and transfer learning (0.76, 0.59, and 0.74 for DOR, MOR, and KOR, respectively) for training. The GCN models perform better on this simpler problem, with excellent recall (0.93, 0.85, 0.80 from direct learning and 0.95, 0.88, 0.89 from transfer learning for DOR, MOR, and KOR, respectively) and precision (0.70, 0.80, 0.90 from direct learning and 0.75, 0.84, 0.87 from transfer learning for DOR, MOR, and KOR, respectively).

**Figure 5.**
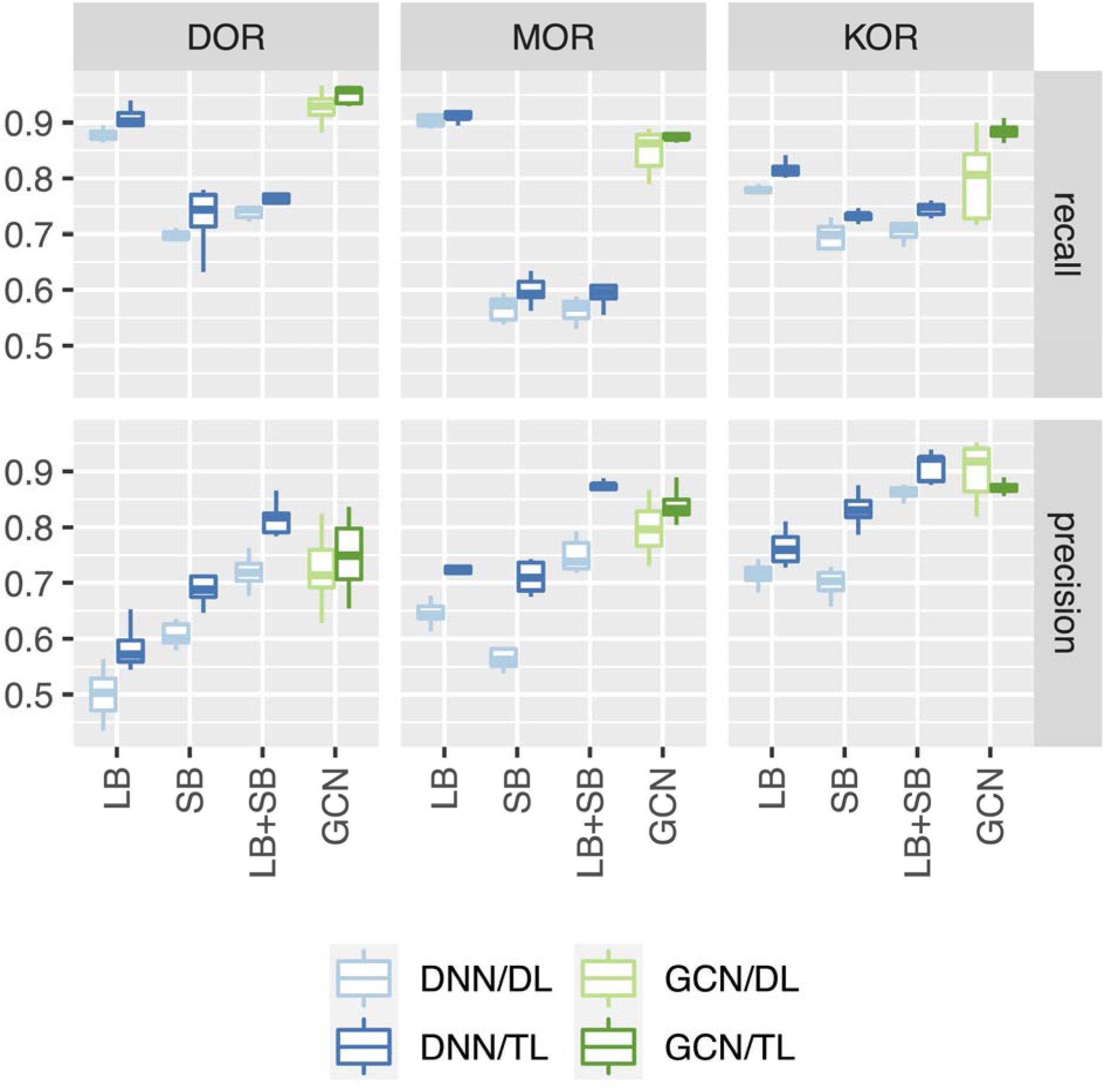
Precision and recall for the prediction of active and inactive molecules from a dataset that includes both inactive and potent active opioid compounds extracted from ChEMBL. The results for the DNN classifiers are shown in light blue for direct learning and dark blue for transfer learning. The results for GCN models are represented in light green for direct learning and dark green and for transfer learning.

## 4. CONCLUSIONS

This study evaluated the effectiveness of integrating LB and SB drug discovery strategies with transfer learning and deep learning models, specifically DNN and GCN classifiers, to enhance the prediction of opioid bioactivity at individual ORs. The research revealed that simple DNN models trained using either LB or SB molecular descriptors exhibited good AUC scores and recall but were not sufficient for practical application due to low precision. The use of combined LB+SB descriptors in the DNN models showed marginal improvements in precision, but notable enhancements were achieved by employing transfer learning and a combination of LB and SB features, improving precision up to 60% on average and increasing recall significantly. This demonstrates that transfer learning and the combined use of LB and SB attributes effectively reduces the false discovery rate and enhances the true positive rate, reducing misclassification and enhancing classification of active ligands.

Comparisons between DNN and GCN classifiers also highlighted that transfer learning provides consistent benefits in model performance. The LB-only GCN architecture outperformed DNN models using LB features in terms of AUC scores, recall, and precision, but DNN models using combined LB+SB descriptors still surpassed the LB-only GCN architecture, particularly in precision.

The study further illustrated that the integration of structural data remains beneficial for predicting ligands with atypical chemical scaffolds that are often misclassified by simpler models. The findings thus affirm the value of integrating transfer learning with hybrid LB+SB-based drug discovery strategies in compound prediction of bioactivities at GPCRs. As more data becomes available in the GPCR field, the performance of deep learning methods is likely to improve, making further exploration and improvement of current AI-based tools both timely and worth pursuing.

## ASSOCIATED CONTENT

Supporting Information

## ACKNOWLEDGMENTS

The authors are grateful to Leslie Salas-Estrada for guidance in the calculation of the structural descriptors. This work was funded by National Institutes of Health grant DA045473. Computations were supported in part through the computational resources and staff expertise provided by Scientific Computing at the Icahn School of Medicine at Mount Sinai, the Clinical and Translational Science Awards (CTSA) grant UL1TR004419 from the National Center for Advancing Translational Sciences, and the Office of Research Infrastructure of the National Institutes of Health under award number S10OD026880. The content is solely the responsibility of the authors and does not necessarily represent the official views of the National Institutes of Health.

