## Supplementary Information for "Enhancing Opioid Bioactivity Predictions through Integration of Ligand-Based and Structure-Based Drug Discovery Strategies with Transfer and Deep Learning Techniques"

Table S1. Composition of the biological activity datasets used for training

Table S2. Composition of the test set of “potent active” opioid ligands from ChEMBL

Table S3. PDB codes of experimental ligand-OR complex structures used to derive SB descriptors

Table S4. Molecular descriptors used for the LB featurization of ligands

Table S5. Performance metrics for DOR DNN models on the test set

Table S6. Performance metrics for MOR DNN models on the test set

Table S7. Performance metrics for KOR DNN models on the test set

Table S8. Performance metrics for GCN models on test set

Table S9. Performance metrics for the DNN and GCN models in classifying potent OR bioactive ligands and inactive ligands from the ChEMBL database.

Figure S1. Hyperparameter optimization for the DNN models using LB features

Figure S2. Hyperparameter optimization for the DNN models using SB features

Figure S3. Hyperparameter optimization for the DNN models using LB+SB features

Figure S4. Hyperparameter optimization for the GCN models

Figure S5. Performance metrics for the DNN classifier on the training set

Figure S6. Salient features for DOR DNN models

Figure S7. Salient features for MOR DNN models

Figure S8. Salient features for KOR DNN models

Figure S9. Representative active ligands that are consistently misclassified by DNN models when trained directly using LB features, but are correctly classified when trained through transfer learning using LB+SB features

Table S1: Composition of the biological activity datasets used for training. The total number of reported agonist and antagonist biological activities (-log_10_ of Ki, IC50, or EC50) at each opioid receptor, excluding linear peptides, was obtained from the IUPHAR/BPS Guide to Pharmacology database. The number of inactive ligands (-log_10_ of Ki, IC50, or EC50 lower than 5) were obtained from ChEMBL. Numbers in parentheses refer to the number of unique chemotypes. The mean and standard deviation of -log_10_ potency values (-log_10_ of Ki, IC50, or EC50) are reported for each group.

|  | | Agonists | | Antagonists | | Inactives | |
| --- | --- | --- | --- | --- | --- | --- | --- |
| **Receptor** | **Number** | | **Potency** | **Number** | **Potency** | **Number** | **Potency** |
| DOR | 40 (36) | | 7.9±1.2 | 30 (21) | 8.1±1.6 | 1672 (1400) | 4.6±0.4 |
| MOR | 54 (47) | | 8.2±1.4 | 33 (29) | 8.9±1.1 | 1201 (1058) | 4.6±0.4 |
| KOR | 68 (48) | | 8.2±1.4 | 33 (23) | 8.5±1.3 | 772 (583) | 4.8±0.3 |
| OR Subfamily | 172 (74) | | 8.1±1.3 | 97 (36) | 8.5±1.4 | 3645 (1936) | 4.6±0.4 |

Table S2: Composition of the test set of “potent active” opioid ligands from ChEMBL. Total number of reported biological activity (-log_10_ of Ki, IC50, or EC50 > 7) at each opioid receptor. Numbers in parentheses refer to the number of unique chemotypes. The mean and standard deviation of -log_10_ potency values (-log_10_ of Ki, IC50, or EC50) are reported for each class.

| **Receptor** | **Number** | **Potency** |
| --- | --- | --- |
| DOR | 1065 (765) | 8.8±0.6 |
| MOR | 1522 (1130) | 8.8±0.7 |
| KOR | 1432 (1017) | 9.1±0.7 |
| OR Subfamily | 4019 (2231) | 8.9±0.7 |

Table S3. Protein Data Bank (PDB) codes of experimental ligand-OR complex structures used to derive SB features. The experimental structural resolution, along with the ligand name and type, and the respective bibliographic references are also reported.

| **Receptor** | **Complex PDB** | **Res. (Å)** | **Ligand PDB** | **Ligand name** | **Ligand type** | **Reference** |
| --- | --- | --- | --- | --- | --- | --- |
| DOR | 6PT3 | 3.3 | OWY | DPI-287 | Agonist | ^28^ |
|  | 4N6H | 1.8 | EJ4 | Naltrindole | Antagonist | ^29^ |
|  | 4EJ4 | 3.4 | EJ4 | Naltrindole | Antagonist | ^30^ |
| MOR | 7UL4 | 2.8 | NG0 | Alvimopan | Antagonist | ^36^ |
|  | 4DKL | 2.8 | BF0 | BF0 | Irreversible antagonist | ^37^ |
|  | 5C1M | 2.07 | VF1 | BU72 | Agonist | ^38^ |
|  | 7U2L | 3.2 | L0X | C5-guano | Bitopic agonist | ^39^ |
|  | 7U2K | 3.3 | KZR | C6-guano | Bitopic agonist | ^39^ |
|  | 8EF5 | 3.3 | 7V7 | Fentanyl | Agonist | ^40^ |
|  | 7SCG | 3 | 8RI | FH210 | G protein-biased/Partial agonist | ^41^ |
|  | 7T2H | 3.2 | EID | Lofentanil | Agonist | ^42^ |
|  | 8EF6 | 3.2 | MOI | morphine | Agonist | ^40^ |
|  | 7T2G | 2.5 | EIG | MP | G protein-biased/Partial agonist | ^42^ |
|  | 7SBF | 2.9 | 8QY | PZM21 | G protein-biased/Partial agonist | ^41^ |
|  | 8EFO | 2.8 | 8QY | PZM21 | G protein-biased/Partial agonist | ^40^ |
|  | 8EFL | 3.2 | WH9 | SR17018 | G protein-biased/Partial agonist | ^40^ |
|  | 8EFB | 3.2 | WH2 | TRV130 | G protein-biased/Partial agonist | ^40^ |
| KOR | 8DZS | 2.65 | U9I | GR89,696 | Agonist | ^31^ |
|  | 8DZR | 2.61 | U9I | GR89,696 | Agonist | ^31^ |
|  | 4DJH | 2.9 | JDC | JDTic | Antagonist | ^32^ |
|  | 6VI4 | 3.3 | JDC | JDTic | Antagonist | ^33^ |
|  | 8DZQ | 2.82 | U99 | momSalB | Full agonist | ^31^ |
|  | 8DZP | 2.71 | U99 | momSalB | Full agonist | ^31^ |
|  | 6B73 | 3.1 | CVV | MP1104 | Agonist | ^34^ |
|  | 7YIT | 3.3 | IVB | Nalfurafine | Full agonist | ^35^ |

Table S4: Molecular descriptors used for the LB featurization of ligands.

| **Variable** | **Description** |
| --- | --- |
| exactmw | Exact molecular weight of the molecule |
| amw | Average molecular weight of the molecule ignoring hydrogens |
| lipinskiHBA | Standard Lipinski HBA definition (number of Ns and Os) |
| lipinskiHBD | Standard Lipinski HBA definition (number of N-H and O-H bonds) |
| NumRotatableBonds | Number of rotatable bonds |
| NumHBD | Number of H-bond donors |
| NumHBA | Number of H-bond acceptors |
| NumHeavyAtoms | Number of heavy atoms |
| NumAtoms | Number of atoms |
| NumHeteroatoms | Number of heteroatoms |
| NumAmideBonds | Number of amide bonds in a molecule |
| FractionCSP3 | Fraction of C atoms that are SP3 hybridized |
| NumRings | Number of rings |
| NumAromaticRings | Number of aromatic rings |
| NumAliphaticRings | Number of aliphatic (containing at least one non-aromatic bond) rings |
| NumSaturatedRings | Number of saturated rings |
| NumHeterocycles | Number of heterocycles |
| NumAromaticHeterocycles | Number of aromatic heterocycles |
| NumSaturatedHeterocycles | Number of saturated heterocycles |
| NumAliphaticHeterocycles | Number of aliphatic (containing at least one non-aromatic bond) heterocycles |
| NumSpiroAtoms | Number of spiro atoms (atoms shared between rings that share exactly one atom) |
| NumBridgeheadAtoms | Number of bridgehead atoms (atoms shared between rings that share at least two bonds) |
| NumAtomStereoCenters | Total number of atomic stereocenters (specified and unspecified) |
| NumUnspecifiedAtomStereoCenters | Number of unspecified stereo atom stereo centers |
| labuteASA | Approximate Surface Area ASA value from Labute (2000) JMGM 18:464 |
| tpsa | Topological polar surface area from Ertl & al. (2000) JMC 43:3714 |
| CrippenClogP | logP value from Wildman & Crippen (1999) JCICS 39:868 |
| CrippenMR | MR value from Wildman & Crippen (1999) JCICS 39:868 |
| chi0v, chi1v, chi2v, chi3v, chi4v | Molecular connectivity indices from Hall & Kier (1991) RCC 2:367-422 |
| chi0n, chi1n, chi2n, chi3n, chi4n | Similar to molecular connectivity, but uses nVal instead of valence |
| hallKierAlpha | The alpha value from Hall & Kier (1991) RCC 2:367-422 |
| kappa1, kappa2, kappa3 | Molecular shape indices from Hall & Kier (1991) RCC 2:367-422 |
| Phi | Flexibility Phi value from Kier (1989) QSAR 8:221 |

**Table S5.** Performance metrics for the DOR DNN models on the test set across 10-fold test splits.

| **Protocol** | **Features** | **Metric** | **antagonist** | **agonist** | **inactive** | **Active Average** |
| --- | --- | --- | --- | --- | --- | --- |
| DL | LB | AUC | 0.93 (0.71,1) | 0.81 (0.67,0.95) | 0.88 (0.77,0.96) | 0.87 (0.82,0.92) |
|  |  | Recall | 0.87 (0.54,1) | 0.63 (0.15,1) | 0.81 (0.8,0.83) | 0.75 (0.58,0.88) |
|  |  | precision | 0.44 (0.25,0.73) | 0.1 (0.04,0.15) | 0.99 (0.98,1) | 0.27 (0.15,0.42) |
|  | SB | AUC | 0.84 (0.52,1) | 0.88 (0.69,0.98) | 0.83 (0.66,0.97) | 0.86 (0.72,0.98) |
|  |  | Recall | 0.83 (0.56,1) | 0.69 (0.36,0.93) | 0.89 (0.84,0.93) | 0.76 (0.63,0.96) |
|  |  | precision | 0.29 (0.15,0.49) | 0.27 (0.14,0.5) | 0.99 (0.97,1) | 0.28 (0.17,0.42) |
|  | LB+SB | AUC | 0.94 (0.81,1) | 0.9 (0.71,0.99) | 0.93 (0.83,0.98) | 0.92 (0.82,0.98) |
|  |  | Recall | 0.86 (0.67,1) | 0.72 (0.36,1) | 0.92 (0.89,0.94) | 0.79 (0.53,0.98) |
|  |  | precision | 0.47 (0.28,0.67) | 0.27 (0.22,0.36) | 0.99 (0.97,1) | 0.37 (0.29,0.45) |
| TL | LB | AUC | 0.96 (0.84,1) | 0.86 (0.74,0.97) | 0.91 (0.85,0.97) | 0.91 (0.87,0.94) |
|  |  | Recall | 0.88 (0.54,1) | 0.76 (0.54,1) | 0.86 (0.83,0.89) | 0.82 (0.75,0.88) |
|  |  | precision | 0.4 (0.19,0.59) | 0.17 (0.12,0.23) | 0.99 (0.99,1) | 0.28 (0.17,0.38) |
|  | SB | AUC | 0.93 (0.71,1) | 0.92 (0.83,0.99) | 0.91 (0.79,0.97) | 0.93 (0.79,0.98) |
|  |  | Recall | 0.82 (0.56,1) | 0.75 (0.36,1) | 0.91 (0.88,0.96) | 0.79 (0.55,0.98) |
|  |  | precision | 0.41 (0.16,0.68) | 0.3 (0.15,0.55) | 0.99 (0.97,1) | 0.36 (0.19,0.57) |
|  | LB+SB | AUC | 0.99 (0.99,1) | 0.96 (0.9,0.99) | 0.97 (0.92,0.99) | 0.98 (0.95,1) |
|  |  | Recall | 0.93 (0.7,1) | 0.77 (0.45,1) | 0.95 (0.93,0.97) | 0.85 (0.65,1) |
|  |  | precision | 0.59 (0.46,0.75) | 0.43 (0.26,0.63) | 0.99 (0.98,1) | 0.51 (0.38,0.61) |

Table S6: Performance metrics for the MOR DNN models on the test set across 10-fold test splits

| **Protocol** | **Features** | **Metric** | **antagonist** | **agonist** | **inactive** | **Active Average** |
| --- | --- | --- | --- | --- | --- | --- |
| DL | LB | AUC | 0.91 (0.75,1) | 0.84 (0.65,0.95) | 0.9 (0.79,0.97) | 0.87 (0.76,0.96) |
|  |  | recall | 0.68 (0,1) | 0.71 (0.44,1) | 0.81 (0.77,0.86) | 0.69 (0.25,0.95) |
|  |  | precision | 0.34 (0,0.59) | 0.2 (0.09,0.3) | 0.98 (0.94,1) | 0.27 (0.07,0.45) |
|  | SB | AUC | 0.85 (0.61,0.98) | 0.84 (0.69,0.94) | 0.84 (0.75,0.92) | 0.85 (0.68,0.95) |
|  |  | recall | 0.8 (0.59,1) | 0.69 (0.56,0.88) | 0.82 (0.77,0.87) | 0.75 (0.58,0.9) |
|  |  | precision | 0.37 (0.19,0.6) | 0.24 (0.18,0.29) | 0.97 (0.96,0.99) | 0.3 (0.21,0.41) |
|  | LB+SB | AUC | 0.94 (0.83,1) | 0.91 (0.81,0.97) | 0.92 (0.84,0.98) | 0.93 (0.86,0.99) |
|  |  | recall | 0.83 (0.56,1) | 0.8 (0.44,1) | 0.9 (0.87,0.92) | 0.82 (0.69,1) |
|  |  | precision | 0.58 (0.21,0.85) | 0.33 (0.22,0.39) | 0.99 (0.97,1) | 0.45 (0.25,0.6) |
| TL | LB | AUC | 0.95 (0.87,1) | 0.87 (0.67,0.97) | 0.93 (0.84,0.98) | 0.91 (0.81,0.98) |
|  |  | recall | 0.78 (0.27,1) | 0.8 (0.5,1) | 0.87 (0.8,0.92) | 0.79 (0.46,1) |
|  |  | precision | 0.46 (0.1,0.78) | 0.31 (0.1,0.51) | 0.99 (0.97,1) | 0.39 (0.15,0.59) |
|  | SB | AUC | 0.96 (0.9,1) | 0.9 (0.77,0.99) | 0.9 (0.79,0.98) | 0.93 (0.85,0.99) |
|  |  | recall | 0.78 (0.5,1) | 0.7 (0.43,0.95) | 0.88 (0.84,0.91) | 0.74 (0.55,0.97) |
|  |  | precision | 0.55 (0.38,0.75) | 0.29 (0.18,0.39) | 0.98 (0.95,1) | 0.42 (0.31,0.56) |
|  | LB+SB | AUC | 0.98 (0.91,1) | 0.98 (0.94,1) | 0.97 (0.91,1) | 0.98 (0.94,1) |
|  |  | recall | 0.92 (0.7,1) | 0.91 (0.65,1) | 0.94 (0.91,0.97) | 0.91 (0.79,1) |
|  |  | precision | 0.8 (0.46,1) | 0.49 (0.36,0.62) | 0.99 (0.98,1) | 0.64 (0.41,0.81) |

Table S7: Performance metrics for the KOR DNN models on the test set across 10-fold test splits

| **Protocol** | **Features** | **Metric** | **antagonist** | **agonist** | **inactive** | **Active Average** |
| --- | --- | --- | --- | --- | --- | --- |
| DL | LB | AUC | 0.96 (0.84,1) | 0.81 (0.61,0.94) | 0.88 (0.74,0.95) | 0.88 (0.8,0.96) |
|  |  | recall | 0.9 (0.67,1) | 0.74 (0.4,1) | 0.79 (0.72,0.83) | 0.82 (0.62,0.98) |
|  |  | precision | 0.49 (0.23,0.76) | 0.28 (0.18,0.42) | 0.98 (0.94,1) | 0.39 (0.23,0.51) |
|  | SB | AUC | 0.93 (0.79,1) | 0.88 (0.8,0.97) | 0.91 (0.86,0.97) | 0.9 (0.81,0.98) |
|  |  | recall | 0.89 (0.75,1) | 0.75 (0.57,1) | 0.76 (0.67,0.86) | 0.82 (0.66,1) |
|  |  | precision | 0.53 (0.42,0.6) | 0.28 (0.2,0.39) | 0.97 (0.93,1) | 0.4 (0.36,0.45) |
|  | LB+SB | AUC | 0.95 (0.78,1) | 0.81 (0.58,0.91) | 0.91 (0.81,0.98) | 0.88 (0.76,0.95) |
|  |  | recall | 0.82 (0.4,1) | 0.64 (0.45,0.83) | 0.88 (0.77,0.96) | 0.73 (0.5,0.91) |
|  |  | precision | 0.66 (0.52,0.83) | 0.43 (0.15,0.7) | 0.97 (0.95,1) | 0.55 (0.37,0.71) |
| TL | LB | AUC | 0.98 (0.94,1) | 0.86 (0.72,0.97) | 0.91 (0.81,0.97) | 0.92 (0.86,0.98) |
|  |  | recall | 0.9 (0.67,1) | 0.75 (0.4,1) | 0.83 (0.77,0.88) | 0.82 (0.65,0.98) |
|  |  | precision | 0.55 (0.34,0.69) | 0.34 (0.2,0.5) | 0.97 (0.94,0.99) | 0.44 (0.31,0.58) |
|  | SB | AUC | 0.99 (0.98,1) | 0.92 (0.86,0.98) | 0.94 (0.91,0.99) | 0.96 (0.92,0.99) |
|  |  | recall | 0.89 (0.75,1) | 0.76 (0.58,0.95) | 0.85 (0.77,0.92) | 0.83 (0.68,0.98) |
|  |  | precision | 0.76 (0.56,1) | 0.36 (0.23,0.48) | 0.97 (0.94,0.99) | 0.56 (0.44,0.65) |
|  | LB+SB | AUC | 1 (0.99,1) | 0.94 (0.91,0.96) | 0.97 (0.95,1) | 0.97 (0.95,0.98) |
|  |  | recall | 0.98 (0.89,1) | 0.77 (0.59,0.88) | 0.94 (0.9,0.99) | 0.88 (0.76,0.94) |
|  |  | precision | 0.87 (0.73,1) | 0.59 (0.44,0.85) | 0.99 (0.96,1) | 0.73 (0.63,0.89) |

Table S8. Performance metrics for the GCN models on the test set across 10-fold test splits.

| **Rec.** | **Prot.** | **Metric** | **antagonist** | **agonist** | **inactive** | **Active Average** |
| --- | --- | --- | --- | --- | --- | --- |
| DOR | DL | ROCAUC | 0.92 (0.64,1) | 0.84 (0.5,0.99) | 0.93 (0.86,0.99) | 0.88 (0.71,0.99) |
|  |  | recall | 0.86 (0.44,1) | 0.71 (0.44,1) | 0.91 (0.87,0.95) | 0.78 (0.65,0.94) |
|  |  | precision | 0.38 (0.23,0.54) | 0.27 (0.09,0.47) | 0.99 (0.98,1) | 0.32 (0.21,0.4) |
|  | TL | ROCAUC | 0.97 (0.84,1) | 0.95 (0.86,0.99) | 0.96 (0.89,0.99) | 0.96 (0.89,0.99) |
|  |  | recall | 0.87 (0.5,1) | 0.81 (0.5,1) | 0.93 (0.9,0.96) | 0.84 (0.69,1) |
|  |  | precision | 0.4 (0.28,0.68) | 0.36 (0.14,0.53) | 0.99 (0.99,1) | 0.38 (0.27,0.49) |
| MOR | DL | ROCAUC | 0.94 (0.82,1) | 0.94 (0.86,0.98) | 0.95 (0.87,0.98) | 0.94 (0.86,0.98) |
|  |  | recall | 0.78 (0.5,1) | 0.82 (0.43,1) | 0.86 (0.76,0.93) | 0.8 (0.53,0.96) |
|  |  | precision | 0.43 (0.2,0.67) | 0.28 (0.11,0.41) | 0.99 (0.99,1) | 0.36 (0.23,0.5) |
|  | TL | ROCAUC | 0.98 (0.93,1) | 0.97 (0.92,1) | 0.98 (0.95,1) | 0.98 (0.93,1) |
|  |  | recall | 0.85 (0.58,1) | 0.94 (0.7,1) | 0.92 (0.89,0.95) | 0.9 (0.69,1) |
|  |  | precision | 0.67 (0.41,1) | 0.43 (0.23,0.63) | 1 (0.99,1) | 0.55 (0.4,0.75) |
| KOR | DL | ROCAUC | 0.99 (0.94,1) | 0.9 (0.73,0.99) | 0.94 (0.88,1) | 0.94 (0.86,0.99) |
|  |  | recall | 0.94 (0.69,1) | 0.77 (0.52,0.94) | 0.9 (0.79,0.98) | 0.86 (0.69,0.97) |
|  |  | precision | 0.64 (0.38,1) | 0.55 (0.23,0.9) | 0.98 (0.96,1) | 0.59 (0.42,0.9) |
|  | TL | ROCAUC | 1 (1,1) | 0.97 (0.92,1) | 0.98 (0.94,1) | 0.98 (0.96,1) |
|  |  | recall | 1 (1,1) | 0.9 (0.7,1) | 0.89 (0.82,0.94) | 0.95 (0.85,1) |
|  |  | precision | 0.79 (0.54,1) | 0.5 (0.23,0.68) | 0.99 (0.97,1) | 0.64 (0.48,0.77) |

Table S9. Performance metrics for the DNN and GCN models in classifying potent OR bioactive ligands and inactive ligands from the ChEMBL database.

| **Rec.** | **features** | **precision DL** | **precision TL** | **recall DL** | **recall TL** |
| --- | --- | --- | --- | --- | --- |
| DOR | LB | 0.5 (0.43,0.55) | 0.58 (0.55,0.64) | 0.88 (0.87,0.89) | 0.9 (0.87,0.94) |
|  | SB | 0.61 (0.58,0.64) | 0.69 (0.65,0.71) | 0.69 (0.64,0.71) | 0.73 (0.65,0.78) |
|  | LB+SB | 0.72 (0.68,0.76) | 0.81 (0.79,0.85) | 0.74 (0.72,0.75) | 0.76 (0.74,0.77) |
|  | GCN | 0.7 (0.51,0.8) | 0.75 (0.66,0.83) | 0.93 (0.9,0.96) | 0.95 (0.93,0.97) |
| MOR | LB | 0.65 (0.61,0.69) | 0.73 (0.69,0.76) | 0.9 (0.89,0.92) | 0.91 (0.9,0.92) |
|  | SB | 0.57 (0.54,0.64) | 0.71 (0.68,0.74) | 0.57 (0.54,0.59) | 0.6 (0.57,0.63) |
|  | LB+SB | 0.75 (0.72,0.79) | 0.87 (0.83,0.88) | 0.56 (0.53,0.59) | 0.59 (0.57,0.61) |
|  | GCN | 0.8 (0.73,0.86) | 0.84 (0.8,0.88) | 0.85 (0.8,0.89) | 0.88 (0.86,0.9) |
| KOR | LB | 0.72 (0.69,0.74) | 0.76 (0.73,0.8) | 0.78 (0.76,0.8) | 0.81 (0.79,0.84) |
|  | SB | 0.71 (0.67,0.78) | 0.83 (0.8,0.87) | 0.7 (0.67,0.72) | 0.73 (0.72,0.74) |
|  | LB+SB | 0.86 (0.85,0.88) | 0.91 (0.88,0.93) | 0.71 (0.68,0.72) | 0.74 (0.7,0.76) |
|  | GCN | 0.9 (0.83,0.95) | 0.87 (0.86,0.89) | 0.8 (0.72,0.88) | 0.89 (0.87,0.92) |

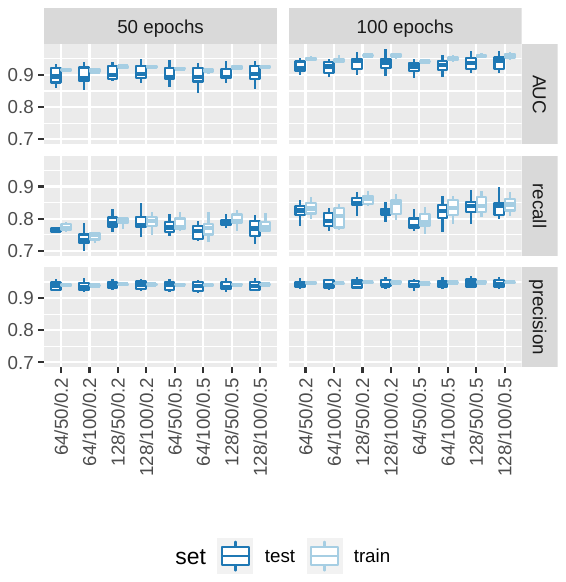

**Figure S1.** Hyperparameter optimization for the DNN models using LB features. The values of the Area under the ROC Curve (AUC), recall, and precision for the train and test sets are provided. The x-axis labels indicate the number of nodes in the hidden layers, the number of nodes in the bypass layer, and the dropout probability. The average AUC, recall, and precision across 10-fold splits for the training set are shown in light blue, while the corresponding values for the test set are depicted in dark blue.

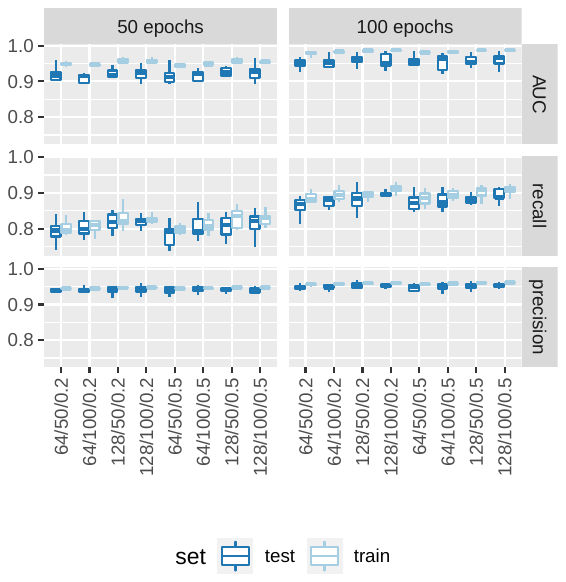

**Figure S2.** Hyperparameter optimization for the DNN models using SB features. The values of the Area under the ROC Curve (AUC), recall, and precision for the train and test sets are provided. The x-axis labels indicate the number of nodes in the hidden layers, the number of nodes in the bypass layer, and the dropout probability. The average AUC, recall, and precision across 10-fold splits for the training set are shown in light blue, while the corresponding values for the test set are depicted in dark blue.

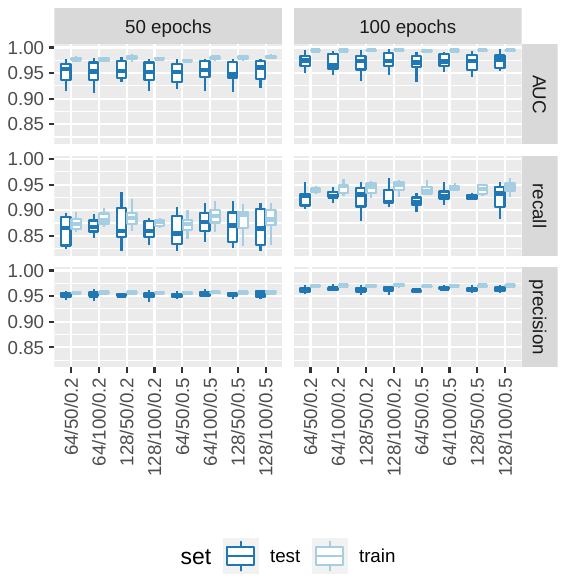

**Figure S3.** Hyperparameter optimization for the DNN models using LB+SB features. The values of the Area under the ROC Curve (AUC), recall, and precision for the train and test sets are provided. The x-axis labels indicate the number of nodes in the hidden layers, the number of nodes in the bypass layer, and the dropout probability. The average AUC, recall, and precision across 10-fold splits for the training set are shown in light blue, while the corresponding values for the test set are depicted in dark blue.

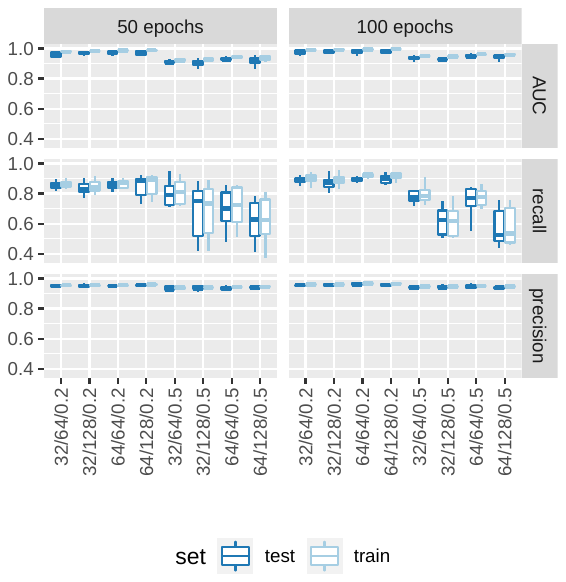

**Figure S4.** Hyperparameter optimization for the GCN model. The values of the Area under the ROC Curve (AUC), recall, and precision for the train and test sets are provided. The x-axis labels indicate the number of nodes in the hidden layers, the number of nodes in the bypass layer, and the dropout probability. The average AUC, recall, and precision across 10-fold splits for the training set are shown in light blue, while the corresponding values for the test set are depicted in dark blue.

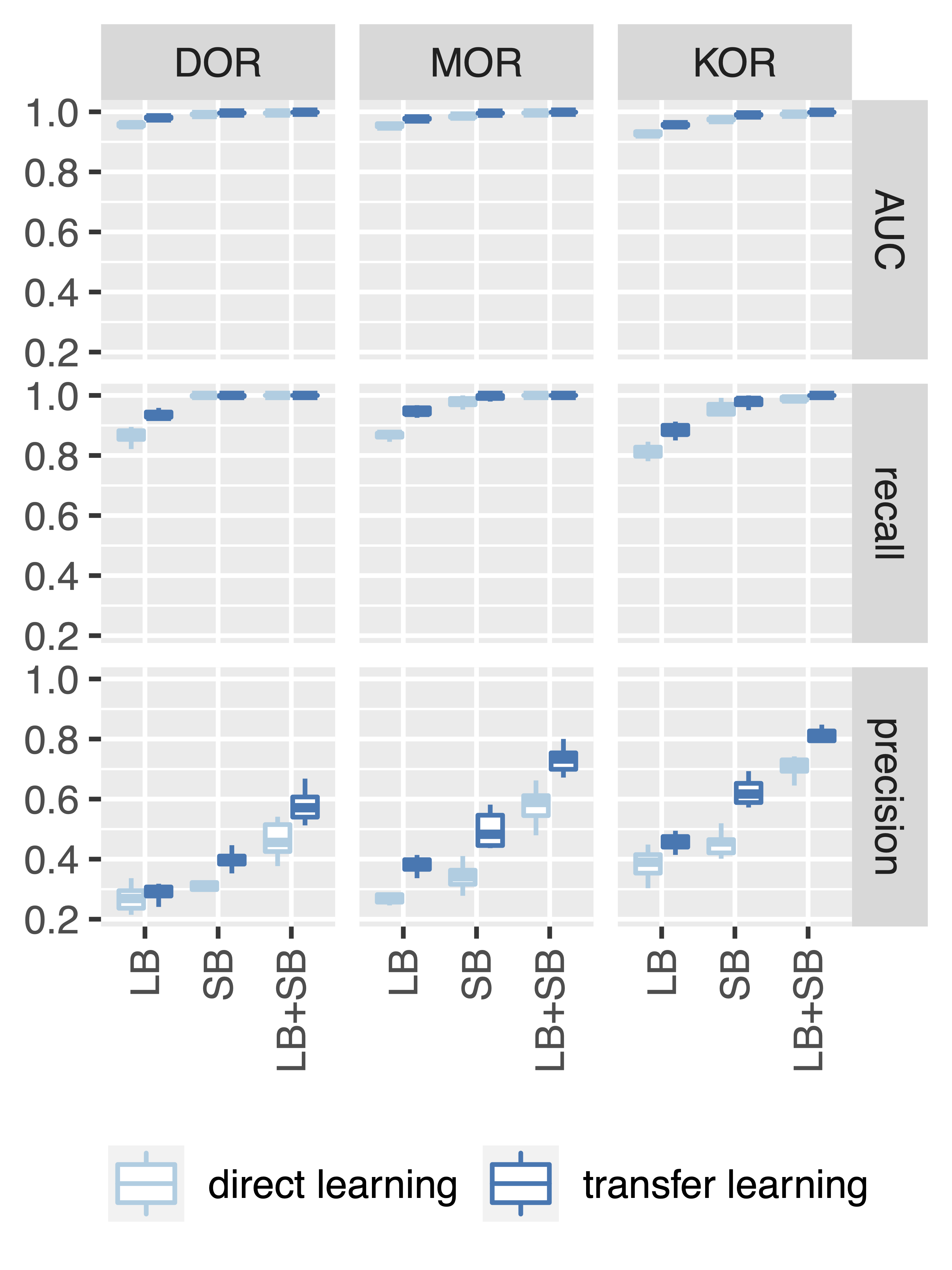

**Figure S5.** Performance metrics for the DNN classifier on the training set across 10-fold training splits.

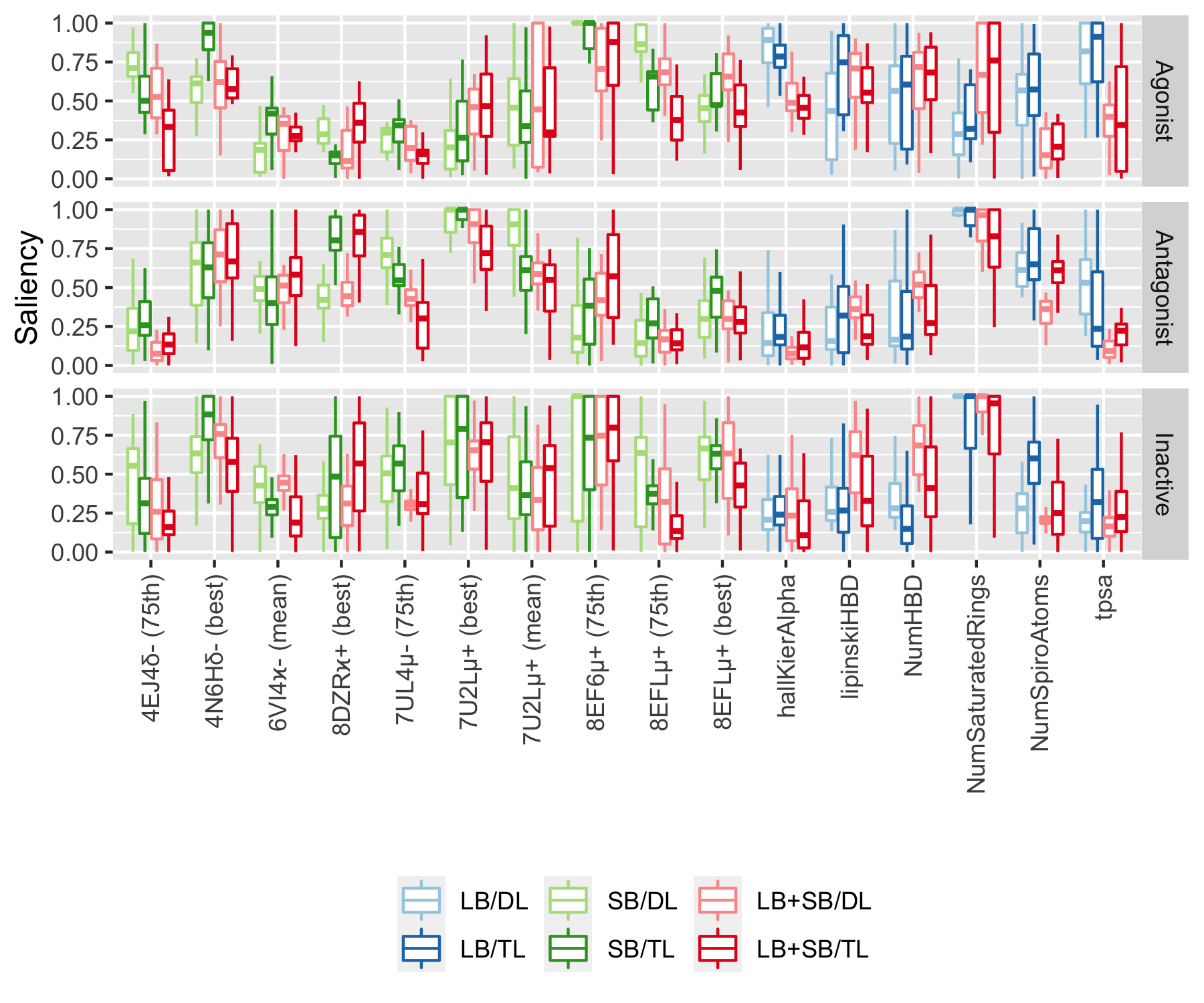

**Figure S6.** Salient features for the DOR DNN models. Only features with an average saliency greater than 0.5 in at least one class are included. Saliency values for the DNN models using LB, SB, and combined LB+SB features are denoted by blue, green, and red, respectively. SB features are labeled by the PDB code, followed by the Greek letter indicating the receptor subtype, and a plus or minus sign indicating whether the ligand in the binding pocket is an agonist or antagonist, respectively. Ligand-based features correspond to the following properties: Wildman & Crippen logP value (CrippenClogP), fraction of cp3 Carbon atoms, Lipinsky’s definition of H-bond donor and acceptor characteristics (lipinskyHBA and lipinskyHBD, respectively), number of H-bond donors and acceptors (NumHBA and NumHBD, respectively), number of amide bonds (NumAmideBonds), number of saturated rings (NumSaturatedRings), and number of spiro atoms, i.e., atoms shared by two rings that share a unique atom (NumSpiroAtoms).

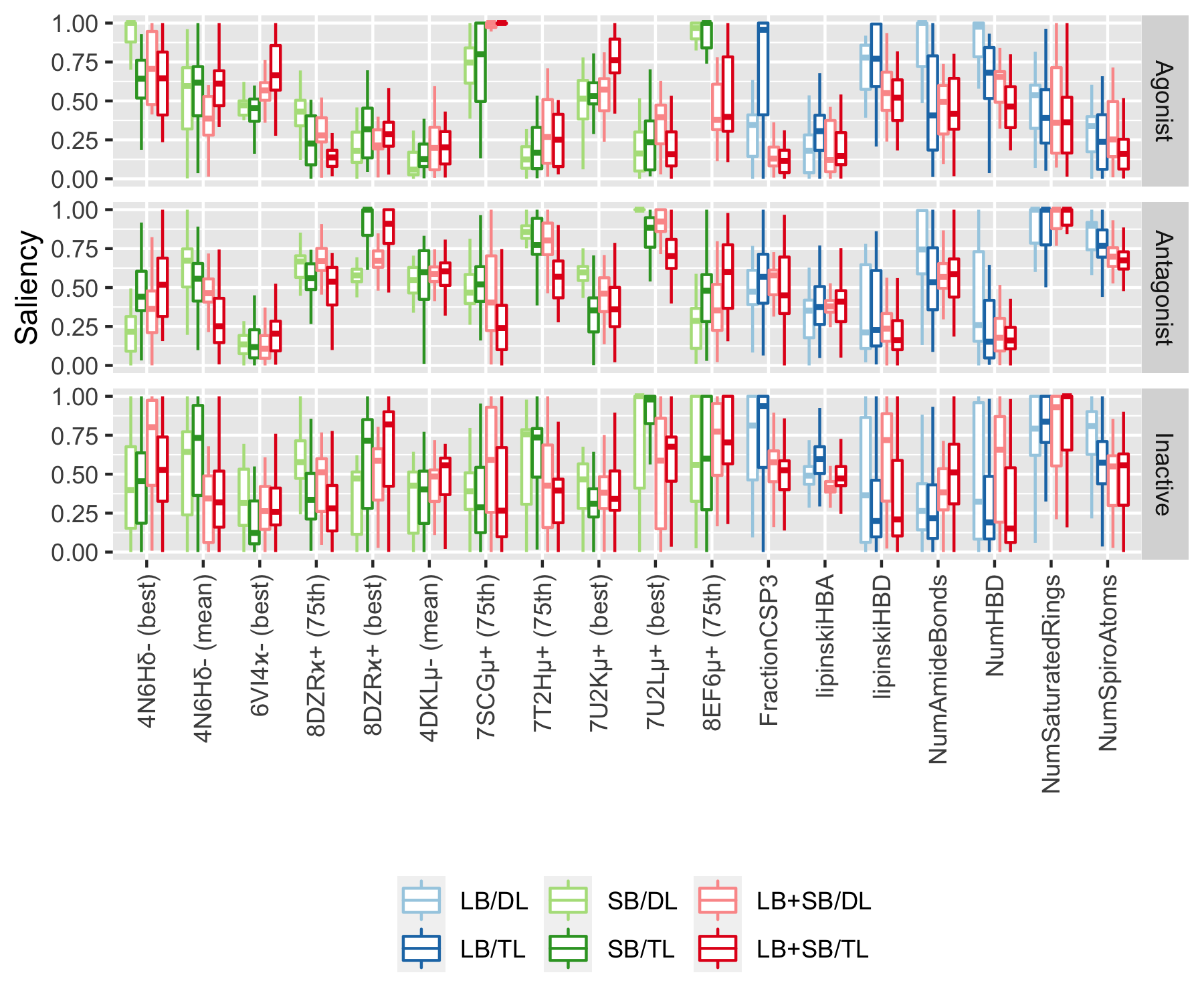

**Figure S7.** Salient features for the MOR DNN models. Only features with an average saliency greater than 0.5 in at least one class are included. Saliency values for the DNN models using LB, SB, and combined LB+SB features are denoted by blue, green, and red, respectively. SB features are labeled by the PDB code, followed by the Greek letter indicating the receptor subtype, and a plus or minus sign indicating whether the ligand in the binding pocket is an agonist or antagonist, respectively. Ligand-based features correspond to the following properties: Wildman & Crippen logP value (CrippenClogP), fraction of cp3 Carbon atoms, Lipinsky’s definition of H-bond donor and acceptor characteristics (lipinskyHBA and lipinskyHBD, respectively), number of H-bond donors and acceptors (NumHBA and NumHBD, respectively), number of amide bonds (NumAmideBonds), number of saturated rings (NumSaturatedRings), and number of spiro atoms, i.e., atoms shared by two rings that share a unique atom (NumSpiroAtoms).

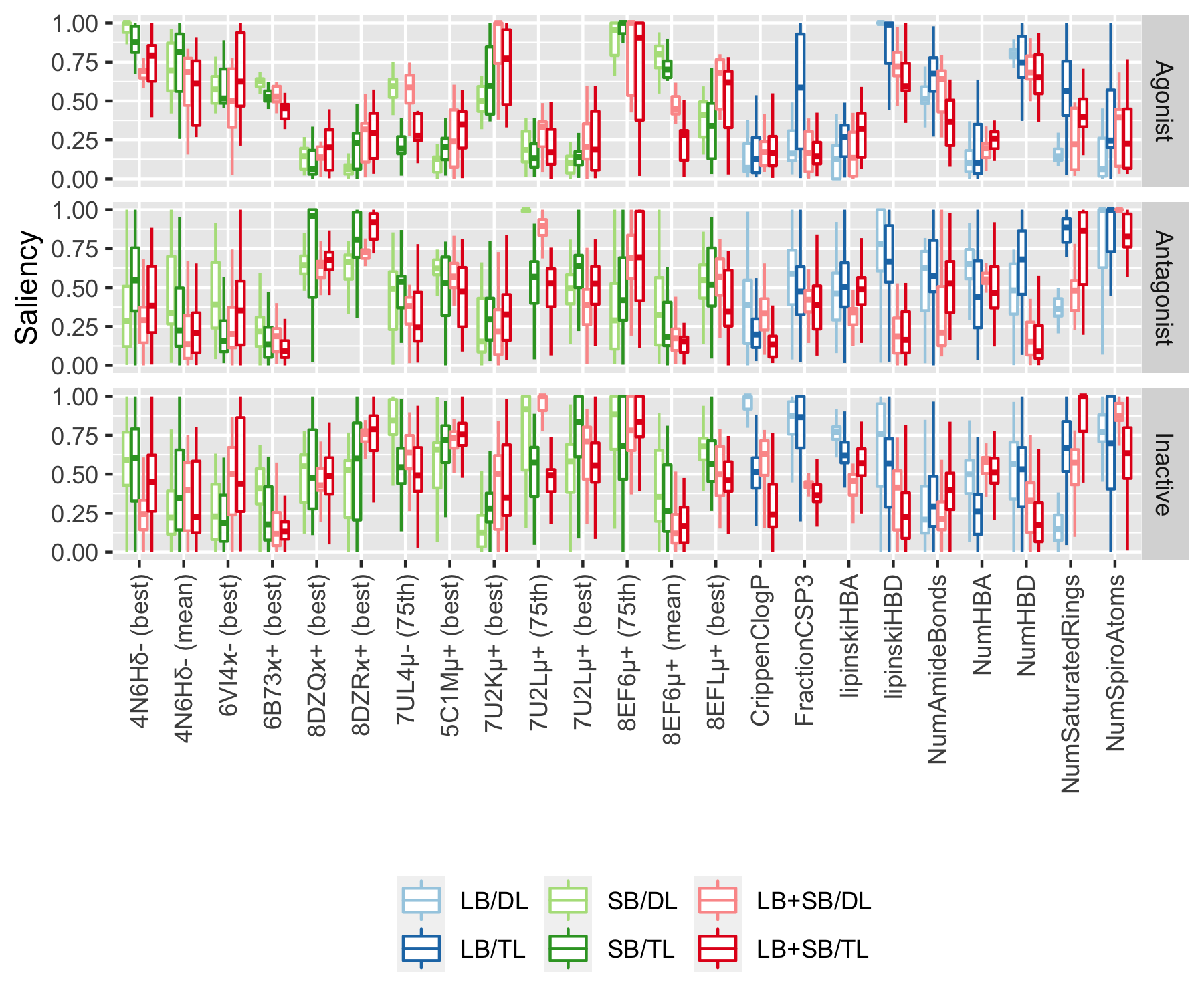

**Figure S8.** Salient features for the KOR DNN models. Only features with an average saliency greater than 0.5 in at least one class are included. Saliency values for the DNN models using LB, SB, and combined LB+SB features are denoted by blue, green, and red, respectively. SB features are labeled by the PDB code, followed by the Greek letter indicating the receptor subtype, and a plus or minus sign indicating whether the ligand in the binding pocket is an agonist or antagonist, respectively. Ligand-based features correspond to the following properties: Wildman & Crippen logP value (CrippenClogP), fraction of cp3 Carbon atoms, Lipinsky’s definition of H-bond donor and acceptor characteristics (lipinskyHBA and lipinskyHBD, respectively), number of H-bond donors and acceptors (NumHBA and NumHBD, respectively), number of amide bonds (NumAmideBonds), number of saturated rings (NumSaturatedRings), and number of spiro atoms, i.e., atoms shared by two rings that share a unique atom (NumSpiroAtoms).

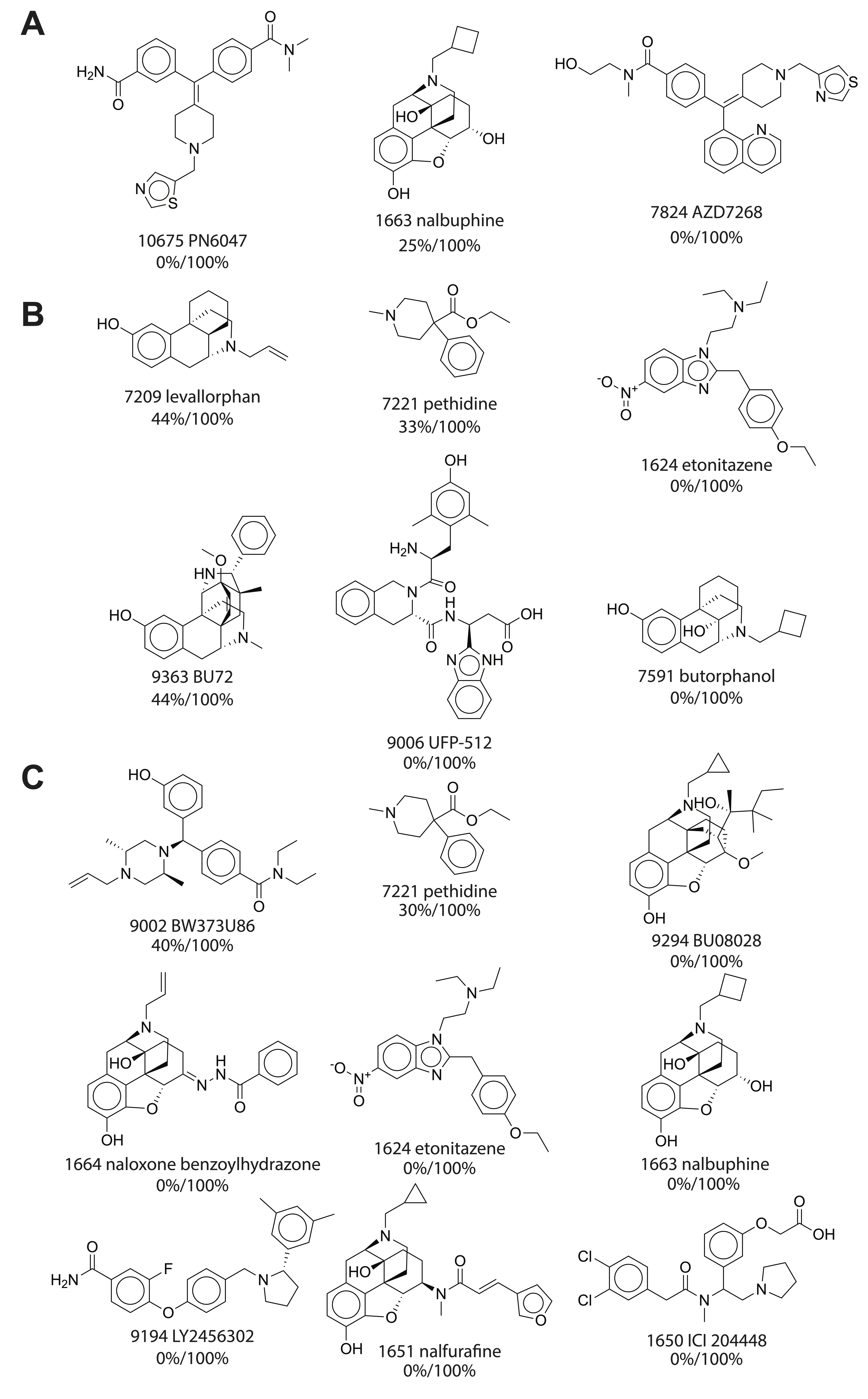

**Figure S9.** Representative active ligands for DOR, MOR, and KOR (in panels A, B, and C, respectively) that are consistently misclassified by DNN models when trained directly using LB features, but are correctly classified when trained through transfer learning using LB+SB features

The fractions indicating how often the ligands are correctly predicted by DNN models trained directly using LB features, as well as by DNN models trained using transfer learning of LB+SB features, are noted beneath the name of each ligand.
